# Word selectivity in high-level visual cortex and reading skill

**DOI:** 10.1101/296061

**Authors:** Emily C. Kubota, Sung Jun Joo, Elizabeth Huber, Jason D. Yeatman

**Affiliations:** Institute for Learning & Brain Sciences; Department of Speech and Hearing Sciences, University of Washington, Seattle, WA, 98195, USA; Department of Psychology, Pusan National University, Busan, Republic of Korea

**Author notes:** These authors contributed equally to this work (Emily C. Kubota, Sung Jun Joo). Correspondence: Emily C. Kubota, Institute for Learning & Brain Sciences Portage Bay Building, Box 357988 University of Washington Seattle, WA 98195, USA, Sung Jun Joo, Department of Psychology, Pusan National University, 2, Busandaehak-ro, 63beon-gil, Geumjeong-gu, Busan, Republic of Korea 46241.

**Keywords:** Visual word form area (VWFA), dyslexia, fusiform face area (FFA), development

## Abstract

Word-selective neural responses in human ventral occipito-temporal cortex (VOTC) emerge as children learn to read, creating a visual word form area (VWFA) in the literate brain. It has been suggested that the VWFA arises through competition between pre-existing selectivity for other stimulus categories, changing the topography of VOTC to support rapid word recognition. Here, we hypothesized that competition between words and objects would be resolved as children acquire reading skill. Using functional magnetic resonance imaging (fMRI), we examined the relationship between responses to words and objects in VOTC in two ways. First, we defined the VWFA using a words > objects contrast and found that only skilled readers had a region that responded more to words than objects. Second, we defined the VWFA using a words > faces contrast and examined selectivity for words over objects in this region. We found that word selectivity strongly correlated with reading skill, suggesting reading skill-dependent tuning for words. Furthermore, we found that low word selectivity in struggling readers was not due to a lack of response to words, but to a high response to objects. Our results suggest that the fine-tuning of word-selective responses in VOTC is a critical component of skilled reading.

## 1. Introduction

Ventral occipito-temporal cortex (VOTC) consists of distributed and overlapping patches of cortex that selectively respond to different categories of images (Grill-Spector & Weiner, 2014). While selectivity for each category—such as faces, places, tools, limbs, and words—has been extensively studied (Bracci, Cavina-Pratesi, Ietswaart, Caramazza, & Peelen, 2012; Dehaene & Cohen, 2011; Downing et al., 2001; Epstein & Kanwisher, 1998; Kanwisher, McDermott, & Chun, 1997; Weiner & Grill-Spector, 2010) we still lack an understanding of how selectivity for these categories emerges in VOTC during development and in relation to learning. Of particular interest, is a region in the left occipito-temporal sulcus, the visual word form area (VWFA), that selectively responds to words compared to other categories of images (Dehaene & Cohen, 2011; McCandliss & Noble, 2003; Petersen, Fox, Posner, Mintun, & Raichle, 1988; Posner, Petersen, Fox, & Raichle, 1988; Wandell, Rauschecker, & Yeatman, 2012). Learning to read plays a critical role in the development of this region, as it is only word-selective in literate as opposed to illiterate adults (Dehaene et al., 2010), and in older children who have received reading instruction as opposed to younger, pre-reading children (Brem et al., 2010; Saygin et al., 2016). Moreover, in children with dyslexia, VOTC is the most consistently reported location of neural deficits (Maisog, Einbinder, Flowers, Turkeltaub, & Eden, 2008; Paulesu, Danelli, & Berlingeri, 2014; Richlan, Kronbichler, & Wimmer, 2011), further emphasizing the importance of this region for skilled reading. It has been suggested that word selectivity in VOTC arises through competition between pre-existing selectivity for other categories of images, changing the topography of VOTC to support rapid word recognition (Dehaene et al., 2010; Dehaene, Cohen, Morais, & Kolinsky, 2015; Dehaene & Cohen, 2007).

How does the process of learning to read change the topography of VOTC to accommodate word-selective cortex? There are many different ways that this process might unfold. Previous literature has focused on competition between words and faces for cortical territory (Dehaene et al., 2010; Monzalvo, Fluss, Billard, Dehaene, & Dehaene-Lambertz, 2012; Plaut & Behrmann, 2011; Yeatman & Norcia, 2016), but there is also evidence suggesting that face selectivity may be stable over development (Kanwisher, 2010; Kuefner, de Heering, Jaques, Palmero-Soler, & Rossion, 2010; McKone, Crookes, Jeffery, & Dilks, 2012). Another possibility is that the VWFA emerges within a general object-selective circuit in VOTC and that, over the course of learning, object responses are pruned away leaving a region that is specialized for words. In line with this hypothesis, it has been argued that both words and objects elicit comparable neural activity in much of VOTC (Kherif, Josse, & Price, 2011; Mano et al., 2013; Price & Devlin, 2003; Wright et al., 2008). Furthermore, the selectivity for words compared to objects in the VWFA differs between adults and children, suggesting a relationship between expertise with text and the relative response to these two image categories in the VWFA (Centanni et al., 2017).

To understand how reading skill shapes tuning properties in the VWFA irrespective of age, we measured selectivity for words over objects in the VWFA of both skilled and struggling readers (i.e., developmental dyslexia). An important point to consider is how methodological differences among studies might affect inferences about VOTC topography: The VWFA is a small patch of cortex that is just a few millimeters away from regions with completely different response patterns, and the location of the VWFA is variable among subjects (Glezer & Riesenhuber, 2013). Using a large smoothing kernel and analyzing data on a standardized template effectively averages the response of the VWFA, fusiform face area (FFA), object- and limb-selective regions. Thus, it is critical to define ROIs in an individual’s native space to examine tuning properties of the VWFA in relation to reading skill. Here, we compared the response to words and objects in ROIs defined in VOTC of individual brains to test whether the VWFA is progressively fine-tuned for words in children with high reading proficiency.

## 2. Methods

### 2.1. Participants

Twenty-four subjects (9 female), ages 7 - 12 years old (M = 9.94, SD = 1.57) participated in this study. Subjects were recruited from the University of Washington Research & Dyslexia Research Database (http://ReadingAndDyslexia.com). All reported normal or corrected-to-normal vision, had an IQ within the normal range (M = 112, SD = 16), were native speakers of English, and had no history of neurological disorder. Twenty-two out of twenty-four were right handed. Prior to their scan, subjects were taken to the MRI simulator in order to acclimate them to the scan environment, and practice holding still. Eleven out of the 24 subjects were diagnosed with dyslexia (based on parent report), however many children who have reading difficulty do not have an official diagnosis of dyslexia, and the criterion for diagnosis is known to vary among practitioners (Siegel, 2006). For the purposes of our analysis, we relied on an in-laboratory reading assessment (rather than parent report) for consistency and divided children into two groups: skilled readers, those with a TOWRE Index >= 85 (n = 8), and struggling readers, those with a TOWRE Index < 85 (n = 16). The cutoff of 85 signifies one standard deviation below the population mean on this aged-normed, standardized measure of reading skill, and is frequently used as a criterion for defining dyslexia in other studies (Rimrodt et al., 2009; B. A. Shaywitz et al., 2002). All procedures, including recruitment, consent, and testing, followed the guidelines of the University of Washington Human Subjects Division and were reviewed and approved by the UW Institutional Review Board.

### 2.2. Reading Measurements

On the same day as their scan, subjects participated in a behavioral session in which they completed a series of behavioral tests. Reading scores were measured using the Test of Word Reading Efficiency (TOWRE-2), which measures the number of sight words (sight word efficiency, SWE) and pseudowords (phonemic decoding efficiency, PDE) read in 45 seconds. They also were assessed using subtests from the Woodcock-Johnson IV (WJ), which measures untimed sight word and pseudoword reading. TOWRE and WJ measures of reading are highly correlated, but also index slightly different aspects of skilled reading. The TOWRE measures the speed and automaticity or word recognition, while the WJ measures the ability to apply orthographic knowledge to decoding difficult words and pseudowords. Subjects also completed the Wechsler Abbreviated Scale of Intelligence (WASI-II) Matrix Reasoning, and Vocabulary subtests.

### 2.3. Functional MRI Data Acquisition

Functional MRI was performed at The University of Washington Diagnostic Imaging Science Center (DISC) on a Philips Achieva 3T scanner. A whole-brain anatomical volume at 0.8×0.8×0.8 mm resolution was acquired using a T1-weighted MPRAGE (magnetization-prepared rapid gradient echo) sequence. Brain tissue was segmented into gray matter, white matter, and CSF with freesurfer (Fischl et al., 2002), and functional data was visualized on the cortical surface. Functional MRI data were acquired using an echo-planar imaging (EPI) sequence (3×3×3 mm voxels, repetition time 2s, echo time 25ms, flip angle 79°, field of view = 240×240 with 36 oblique slices prescribed parallel to the ventral surface).

### 2.4. Functional MRI Stimuli and Task

Figure 1 shows examples of the stimuli used in the experiment. The stimuli come from the fLoc functional localizer package (Stigliani, Weiner, & Grill-Spector, 2015). The details of the stimuli are described in Stigliani et al. (2015). Briefly, subjects were shown images of text (pseudowords), objects (cars and instruments) and faces (child and adult faces), which were embedded in a phase-scrambled noise pattern. Each phase scrambled patch covered 20 degrees of visual angle. This stimulus set was designed to control for the low level properties of the images (e.g. luminance and contrast), while maintaining clear image categories, in order to measure VOTC tuning to image category without the confounds of overlearned stimulus properties (e.g. courier font). Thus the text was rendered at various oblique angles, with texture added to the letters, and random positions around fovea. Even though there are still differences in some low-level image properties, this stimulus set allows us to study a more abstract representation of image category than had we used black text rendered on a white background. Stimuli were presented in a block design experiment, and each block consisted of eight images presented for 500ms each (400ms stimulus duration + 100ms inter-stimulus interval), for a total of four seconds per block (Figure 1D). There were ten blocks per each stimulus category, plus eleven blank blocks (baseline) and the block order was randomized in each scan run. Subjects were asked to press a button every time an image repeated (one-back task). A repetition occurred in one third of blocks (~4% of stimulus presentations) to minimize potential motion due to button press. Subjects completed two scan runs. A blocked design experiment was used to maximize signal to noise ratio (SNR). However, the limitation of this approach is that we are not able to examine differences in neural response while controlling for performance effects across age (Brown et al., 2005), or reading skill.

**Figure 1.**
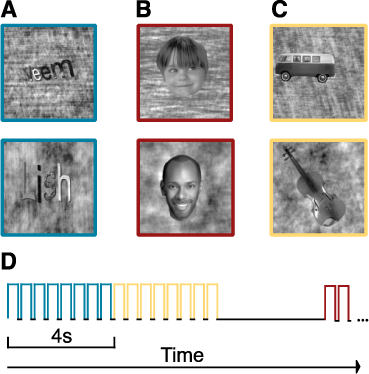
Example stimuli for each stimulus category. (A) Words. Four-letter pseudowords were presented during the word block. (B) Faces. Children and adults’ faces were shown during the face block. (C) Objects. Cars and musical instruments were displayed during the object block. (D) Timing of the functional localizer. In each categorical block, eight images were presented for 400ms followed by 100ms of fixation. Blank blocks (4s of fixation) were randomly interleaved throughout the experiment. During the scan, subjects were instructed to fixate at the fixation mark in the center of display and performed a one-back task.

### 2.5 Data Analysis

Functional MRI data were analyzed using Vistasoft (https://github.com/vistalab/vistasoft). GLMdenoise (Kay, Rokem, Winawer, Dougherty, & Wandell, 2013) was used to improve the signal-to-noise ratio (SNR) of the data before conducting a general linear model (GLM) analysis in Vistasoft. GLMdenoise uses PCA to identify noise sources that are then removed from the time-series using a GLM. Cross-validation is used to determine the optimal number of noise sources to remove from the data without removing any task-related signal. Three subjects with excessive motion (> 2 voxels) were excluded from our analysis. One subject was excluded for having no functional region of interests (including FFA), which may suggest poor data quality or lack of attention to the fMRI task. Twenty subjects (8 female, 12 male; 19 right handed, 1 left handed; 8 skilled readers, 12 struggling readers) were included in subsequent analyses.

Functional regions of interest (ROIs) were defined in individual subjects’ native space. The visual word form area (VWFA) is defined as voxels in the lateral fusiform, occipitotemporal sulcus and inferior temporal gyrus, that selectively respond to words compared to other stimuli (Yeatman, Rauschecker, & Wandell, 2013). However, the literature is inconsistent in which comparison categories are used as a baseline to define the VWFA with different studies using checkerboards (Cohen et al., 2002; Szwed et al., 2011; Yeatman et al., 2013), phase scrambled words (Glezer & Riesenhuber, 2013; Yeatman et al., 2013), objects (Grill-Spector & Weiner, 2014), or fixation (Ben-Shachar, Dougherty, Deutsch, & Wandell, 2011; Boros et al., 2016). For each subject we defined word-selective regions meeting the anatomical criterion of the VWFA using two different contrasts: (1) words vs. objects (VWFA_obj_) and (2) words vs. faces (VWFA_face_). These regions were overlapping for all the subjects that had both regions, which is consistent with the expectation that different contrasts will identify the same region in literate adults. Since we are interested in comparing the relationship between word and object response in word-selective cortex, these regions allow us to examine this relationship in two ways. Our VWFA_obj_ is the result of a direct comparison between word response and object response, and allows us to establish the existence of a region that selectively responds to words over objects in skilled readers. Our VWFA_face_ allows us to index individual differences in word selectivity in a region that is defined independently of object response.

For all ROIs we selected voxels in VOTC that meet our criterion, a threshold of p < 0.001 (uncorrected), with the following exceptions. For those who we could not define VWFA_obj_ (n = 10) or VWFA_face_ (n = 4) using this threshold we tested to see whether we could define these ROIs at a more lenient threshold (p < 0.01). We remained unable to define VWFA_obj_ region using the lenient threshold in all ten subjects who did not have VWFA_obj_; this finding confirms that the subjects without VWFA_obj_ do not have a region responding to words compared to objects even at a more lenient threshold. We were, however, able to define VWFA_face_ in an additional two of the four subjects without VWFA_face_ at this more lenient threshold, and these two subjects are included in subsequent analyses of word selectivity in VWFA_face_. VWFA_face_ in these subjects was located in the correct anatomical location. Moreover, we confirmed that excluding these two subjects from the main analyses did not change the pattern of results. For one subject the VWFAface was highly right lateralized so we used this right hemisphere region as it is known that VWFA is right lateralized in a small subset of individuals (Cohen et al., 2002). Fusiform face area (FFA) ROIs were defined using a face vs. object contrast for 15/20 of the subjects. For the remaining 5/20 subjects (three struggling readers, and two skilled readers) who we could not localize a FFA using a face vs. object contrast, we used a face vs. baseline contrast and selected voxels meeting the anatomical criterion for the FFA.

Within the VWFA_face_ region, we calculated a word selectivity index (SI) as follows,

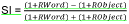

where *RWord* is the BOLD response for words and *RObject* is the BOLD response for objects. We added 1 to the BOLD response for each stimulus category to avoid the SI bounding at one when *RObjectis* negative, in which the index does not properly represents the relationship between the two measurements. This effectively decreases the index by half while maintaining the relationship between the two measurements. A positive selectivity index denotes a greater response to words compared to objects, whereas a negative selectivity index denotes a greater response to objects compared to words. A selectivity index of zero, would denote equivalent response to words and objects.

In order to ensure that there was no difference in data quality among our subjects, we defined a stimulus-responsive region in early visual cortex in 20/20 subjects using a (face + word + object > fixation) contrast at a threshold of p < 0.0000000001 (uncorrected). We found that there was no correlation between variance explained (R^2^ in the GLM) in the early visual cortex response and reading skill (*r* = 0.28, *p* = 0.23). In addition, there was no difference in BOLD responses to words (*t*(18) = −0.04, *p* = 0.97), faces (*t*(18) = 0.18, *p* = 0.86), or objects (*t*(18) = −0.43, *p* = 0.67) in skilled versus struggling readers in early visual cortex. These analyses indicate that there was no difference in data quality or compliance between the groups.

## 3. Results

### 3.1. Skilled readers have a VWFA that responds selectively to words compared to objects and faces

In the literate adult brain the VWFA is more selective for words compared to objects. Moreover, it is interdigitated, and partially overlapping with object-selective regions and is lateral and non-overlapping with face selective regions (Grill-Spector & Weiner, 2014). These observations suggest that proficient readers should have a region that responds more to words compared to objects and words compared to faces.

In 10 out of 20 children we could localize a VWFA_obj_ in VOTC, and as expected, the subjects for whom we could localize this region were significantly stronger readers than the subjects who did not have a VWFA_obj_. Reading skills measured by the TOWRE Index, WJ Basic Reading Skills Composite (standardized scores with mean of 100 and the standard deviation of 15), as well as TOWRE Sight Word Efficiency (raw score, number of sight words read in 45 seconds) were higher for the subjects with a VWFA_obj_ compared to the subjects without a VWFA_obj_ (Figure 2A; TOWRE Index, *t*(18) = 6.13, *p* = 0.00001; WJ BRS, *t* (18) = 5.04, *p* = 0.0001; TOWRE SWE, *t* (18) = 7.21, *p* = 0.000001).

**Figure 2.**
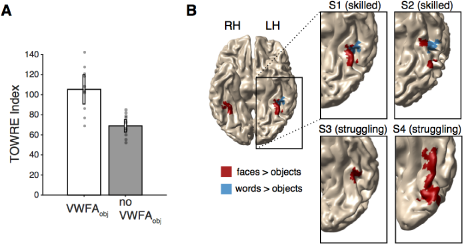
Only skilled readers have a region responding to words over objects. (A) Reading skill in subjects with and without VWFA_obj_. Reading skill is indexed by the TOWRE index (standard score; M = 100, SD = 15). Y-axis represents the mean TOWRE index within each group. Subjects with a VWFA_obj_ (n = 10) are significantly stronger readers than those without a VWFA_obj_ (n = 10) (p = .00001). The gray data points show each individual’s TOWRE index. The error bars indicate the SEM across subjects. (B) VWFA_obj_ (blue) and face-object (red) ROIs in example subjects. The top and bottom rows show example skilled and struggling readers, respectively. RH: right hemisphere, LH: left hemisphere.

To ensure that this finding didn’t reflect differences in data quality due to head motion, we tested whether there was a difference in head motion between subjects with VWFA_obj_ and without VWFA_obj_and found that there was no difference (*t*(18) = −1.27, *p* = .22; homogeneity of variance was confirmed with an O’Brien test F(1,18) = 0.07, *p* = .80 (O’Brien, 1979)). Figure 2B shows VWFA_obj_ (blue) and FFA (red) regions in four example subjects. The top row shows data for two skilled readers and the bottom row shows two struggling readers. These findings suggest that word selectivity in VOTC only emerges after acquiring a high level of reading proficiency and that struggling readers do not have a region that selectively responds to words compared to objects.

In all ten subjects for whom we could define a VWFA_obj_, we could also define a VWFA_face_. Critically, in all of these subjects the VWFA_obj_ and the VWFA_face_ ROIs were overlapping, indicating that both contrasts are localizing the same word-selective region for the skilled readers (see Figure 3A; VWFA_obj_ in blue, VWFA_face_ in dashed outline). Next, we tested whether we could find a VWFA_face_ in the subjects for whom we could not find a VWFA_obj_. In 8 out of 10 children who did not have a VWFA_obj_, we could find a VWFA_face_ (Figure 3A). This finding suggests that VOTC of struggling readers still responds selectively to words compared to faces despite the lack of selectivity over objects. This finding is consistent with the hypothesis that for struggling readers words are processed by a general region that responds equivalently to various types of objects (including words), but that struggling readers do not have a specialized region for word recognition within this general object-selective circuit.

**Figure 3.**
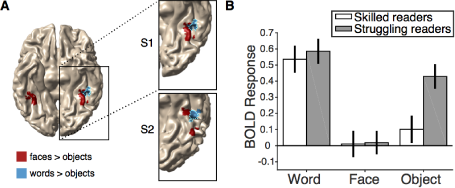
Responses in the VWFA_face_. (A) The FFA is shown in red, VWFA_obj_ is shown in blue, and VWFA_face_ is outlined in cyan for two example, skilled reading, subjects. Note that the two ROI definitions are overlapping indicating that in skilled readers both contrasts identify the same word-selective patch of cortex. (B) BOLD responses (beta weights from the GLM) for each stimulus category (word, face, and object) in skilled (white) and struggling (gray) readers. Skilled readers show word-selective responses (words > objects) while struggling readers show very weak word selectivity in this region with a comparable response to words and objects. The error bars represents the SEM across subjects.

### 3.2. Underactivation versus lack of selectivity in VOTC of struggling readers

Given that previous studies have suggested that VOTC of struggling readers is underactivated during the word presentation (Maisog et al., 2008; Eraldo Paulesu et al., 2014; Richlan et al., 2011; B. A. Shaywitz et al., 2002; S. E. Shaywitz et al., 1998), it seems surprising that we were able to define a region in VOTC using a word versus face contrast in most of our struggling readers. Does VOTC of our struggling readers respond to words strongly at least compared to faces but not selectively compared to objects?

To test this possibility, we divided the 18 subjects with a VWFA_face_ ROI into two groups based on TOWRE index scores (skilled readers, TOWRE index >= 85, n = 8; struggling readers, TOWRE index < 85, n = 10) and assessed the response to each stimulus category in the VWFA_face_ ROI (Figure 3A). Figure 3B shows the beta estimates for each stimulus category (words, faces, and objects) and subject group (skilled and struggling readers). We conducted a mixed-design ANOVA (within-factor (stimulus category) and between-factor (group)). For our statistical analysis, we only compared words and objects because the ROIs were defined using a words > faces contrast, making any comparisons between words and faces redundant. Importantly, there was no main effect of group (F(1,16) = 3.07, *p* = 0.10) indicating that the amplitude of the BOLD response was not lower in struggling compared to skilled readers. There was a main effect of stimulus category, (F(1,16) = 50.04, *p* = .000002) and a significant group by stimulus category interaction (F(1,16) = 12.25, *p* = 0.003). This interaction reflected differences in the relative tuning of the VWFA to different stimulus categories between skilled versus struggling readers. Critically, we found that skilled readers and struggling readers showed an equivalent BOLD response to words (*t*(16) = 0.43, *p* = 0.68). Thus, when this region of cortex is localized within each individual’s brain, there is no evidence of underactivation to words in the struggling readers. In contrast, responses to objects were higher in struggling readers compared to skilled readers (*t*(16) = 2.88, *p* = 0.01). When we conduct the same analysis using a median split (rather than a cutoff of TOWRE < 85), we confirm the same result: There was no main effect of group F(1,16) = 3.38, p = 0.09, there was a main effect of category F (1, 16) = 52.8, *p* = 0.000002, and group by category interaction F(1,16) = 13.8, *p* = 0.002. Conducting an equivalent statistical analysis for the left FFA we do not find a significant group by stimulus category interaction (F(1,16) = .44, *p* = .52) ruling out the possibility that this is a general phenomenon in left VOTC as oppose to an effect specifically within the VWFA.

In summary, we found no evidence supporting underactivation in VOTC of struggling readers. Rather, within individuals, the response to words was greater than objects in skilled readers (*t(7)* = 10.07, *p* = .00002), whereas the same comparison yielded only a marginal effect in struggling readers (*t*(9) = 2.52, *p* = .03). This finding suggests that the VWFA in skilled readers is more finely tuned to words than in struggling readers.

### 3.3. Word selectivity in VOTC predicts reading skill

We have shown that there is a region in VOTC that is selective for words over faces and objects in skilled readers but not in struggling readers. To test directly whether word selectivity is proportional to reading proficiency, we defined a word selectivity index (see Materials and Methods) and assessed the relationship between each individual’s selectivity index and reading skill.

We found that there was a strong correlation between selectivity index in the VWFA_face_ and the TOWRE Index (figure 4A; *r* = 0.71, *p* = 0.001). The TOWRE Index is a good measure to assess an individual’s relative reading ability compared to their peers, however, it does not index absolute reading proficiency because the TOWRE Index would be lower for an older child compared to a younger child if both children read at the same rate. In this sense, the raw score would be a better indication of the relationship between absolute reading skill and selectivity. Indeed, we also found a strong correlation between selectivity index and TOWRE sight word efficiency (figure 4B; *r* = 0.81, *p* = 0.00004). Note that our reading measures (TOWRE Index, TOWRE SWE, WJ BRS) were all correlated with each other, and that our standard reading measures (TOWRE Index, WJ BRS) were not correlated with age (see Table 1). We found that there is a marginal correlation between TOWRE SWE and age (*r* = 0.43, *p* = 0.07) and selectivity and age (figure 4C; *r* = 0.44, *p* = .07), and these correlations might be significant in a larger sample. We used a linear model to assess the contributions of reading skill and age to VWFA selectivity and found significant independent contributions of both variables: TOWRE Index (β = 0.003, SE = 0.0005, p = 0.0004), age (β = 0.002, SE = 0.0007, p = 0.03). This finding demonstrates that both age, and reading skill contribute to word selectivity in our sample, which may explain why TOWRE SWE, a measure that reflects absolute reading ability and is affected both by age and standard reading level, is the strongest correlate of selectivity. We confirmed that there was no correlation between head motion and selectivity index (*r* = −0.18, *p* = 0.47) or head motion and reading score (*r* = −0.36, *p* = .14), ruling out the differences in data quality as a potential confound.

**Figure 4.**
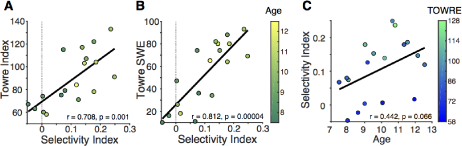
Word selectivity predicts reading skill. (A-B) The correlation between selectivity index and reading skill. Selectivity index is strongly correlated with (A) TOWRE Index (standardized score; M = 100, SD = 15) and (B) TOWRE SWE (raw score). The color bar represents age, in years. (C) Selectivity index and age are marginally correlated. The color bar indicates TOWRE Index score.

**Table 1.** Correlation coefficients for VWFA selectivity, age, TOWRE Index, and TOWRE SWE. ** p < 0.01, *** p < 0.0001

|  | Age | TOWRE | SWE | WJ BRS |
| --- | --- | --- | --- | --- |
| Selectivity | 0.44 | 0.71** | 0.81*** | 0.64** |
| Age |  | 0.13 | 0.43 | 0.06 |
| TOWRE |  |  | 0.90*** | 0.92*** |
| SWE |  |  |  | 0.82*** |

\*\* $p < 0.01$ , \*\*\* $p < 0.0001$

### 3.4. The reading circuitry beyond the VWFA

Reading involves a network of regions, and as such, our word vs. face contrast localized other reading related regions in addition to the VWFA. For example, we were able to localize a response to words in left superior temporal cortex (STC) in 8/8 of our skilled readers, and 7/12 of our struggling readers at a threshold of p < .001 (Supplemental Figure 1A). Those who had an STC response to words were significantly better readers than those who did not (Supplemental Figure 1B; *t*(18) = 2.66, *p* = 0.01).

Unlike in the VWFA, there was no response to visual categories other than words within STC. We performed the same analysis of BOLD responses in left STC, using a mixed-design ANOVA with the within-factor of stimulus category (words, objects) and the between-factor of group (skilled readers, struggling readers). There was a main effect of category F(1,13) = 92.02, p = .0000003, reflecting a higher response to words compared to objects and, in fact, the object response was not significantly above baseline (*t*(14) = −1.15, *p* = 0.27),. This finding makes sense, given that STC is a classic language region and, therefore, is only response to visual stimuli that are linguistic in nature (i.e., words). We found that there was no main effect of group F(1,13) = 0, *p* = 0.99, which demonstrates that there was no difference in overall amplitude of BOLD response in STC between skilled and struggling readers. There was also no group by category interaction F(1,13) = 0.003, *p* = 0.95 (Supplemental Figure 1C). This finding suggests that in both skilled and struggling readers STC responds to words and not to objects, whereas VWFA responds to both words and objects, but to varying degrees depending on reading skill. Of course, it is important to note that we were only able to define STC in a little over half of our struggling readers meaning that many struggling readers show no STC response to words. When we calculate our selectivity index, looking at the ratio of response to words compared to response to objects in STC, we find that there is no correlation with reading skill r = −0.05, p = 0.86 (supplemental figure 1D).

## 4. Discussion

We have demonstrated that word selectivity in VOTC strongly correlates with reading skill and the lack of word selectivity in struggling readers is not due to underactivation in VOTC but from comparable responses to both words and objects. These findings lend support to the idea that there is competition between words and other visual categories for territory in VOTC, and that word selectivity emerges through reorganization in high-level visual cortex during the process of learning to read.

Previous studies have emphasized an underactivation in VOTC to words as a hallmark of dyslexia (Boros et al., 2016; Brunswick, McCrory, Price, Frith, & Frith, 1999; Maisog et al., 2008; E. Paulesu et al., 2001; B. A. Shaywitz et al., 2002; S. E. Shaywitz et al., 1998). Here, we argue that atypical VOTC response in dyslexics is not due to a lack of response to words, but rather a lack of word selectivity. In our struggling readers, there was comparable response to words and objects in VWFA_face_, whereas our skilled readers demonstrated selectivity in which there was a greater response to words than objects. One potential explanation for these seemingly discrepant findings might be the effects induced by the cognitive task subjects perform while viewing words. Most of these previous studies used tasks that target aspects of reading such as reading out loud, rhyming, and lexical decisions. Brunwick et al. (1999) found underactivation in dyslexics compared to controls in their explicit and implicit reading tasks. On the other hand, Shaywitz et al. (1998) used hierarchical tasks (case judgement, single letter rhyme judgement, pseudoword judgement, and semantic comparison) to engage differing degrees of language processing. They found that there was no difference in VOTC between dyslexics and controls for the case judgment task, but there was an increasing difference between dyslexics and controls as the linguistic demands of the task increased, such that the greatest difference between dyslexics and controls was in the semantic comparison task. As subjects perform these tasks, the BOLD response in VOTC will reflect a combination of bottom-up and top-down signals (Kay & Yeatman, 2017). Thus, there are multiple levels of processing in which dyslexics might have a deficit. Previous work has demonstrated that sensitivity to word visibility is correlated with reading skill (Ben-Shachar et al., 2011), and changes with learning, providing evidence for reading skill related differences in bottom-up processing of words. Here, comparing responses to different image categories in a task with minimal linguistic demands, we found differences in tuning properties between skilled and struggling readers. In more linguistically demanding tasks such as rhyme judgement paradigms, there is an additional element of phonological processing that may elucidate additional top-down deficits. The contribution of bottom-up and top-down effects to neural processing deficits in dyslexia is an important point to resolve in future work.

Previous work has proposed that the process of learning to read results in competition between response to words and other categories of visual images (Dehaene & Cohen, 2007; Dehaene et al., 2015). Among various categories, the relationship between words and faces have been extensively studied with regard to the emergence of word selectivity. This research has been motivated by the fact that the locations of word-selective and face-selective areas are in close proximity and face selectivity is right-lateralized in literate adults (Dehaene & Cohen, 2011). It has been shown that an increased response to words after acquiring literacy seems to result in a decreasing response to faces in left VOTC (Dehaene et al., 2010). Both fMRI and ERP responses are more right-lateralized as literacy increases (Dehaene et al., 2010; Pegado et al., 2014). Moreover, decreasing responses to faces predicts higher task performance associated with symbol processing at age of four (Cantlon, Pinel, Dehaene, & Pelphrey, 2011). Recently, a computational population receptive field analysis suggested that face selectivity and character selectivity might undergo competition for foveal coverage (Gomez, Natu, Jeska, Barnett, & Grill-Spector, 2017).

However, there is also evidence showing face processing may be stable over the course of development. Selectivity for faces over other objects emerges very early in development. Four- to six-month old infants show face-selective steady-state visually evoked potentials (Farzin, Hou, & Norcia, 2012) and BOLD responses (Deen et al., 2017), suggesting a rapid emergence of face selectivity in VOTC after less than six-months of experience with visual exposure to faces. Selectivity for faces over other objects measured at four years of age does not seem to change over the course of development (Kuefner et al., 2010), consistent with evidence showing early maturation of face processing measured by numerous behavioral tasks (reviewed in McKone, Crookes, Jeffery, & Dilks, 2012). Furthermore, the right hemisphere advantage of face processing is found in both infants (de Schonen & Mathivet, 1990) and young children (Marcel & Rajan, 1975; Young & Bion, 1980; Young & Ellis, 1985). In the present study, were able to define a region that responds to words greater than faces in the majority of our subjects, whereas we were only able to define a region that responds to words greater than objects in stronger readers. An interesting hypothesis that emerges from our work is that there are multiple stages of learning, where selectivity for words compared to faces emerges in VOTC before object responses are pruned away. Thus, longitudinal and intervention research is warranted to understand developmental trajectories of competition between words and faces given early development of face selectivity in VOTC.

## 5. Conclusions

Previous cross-sectional studies of children have used group comparisons, spatial smoothing, and templates that may make it difficult to observe subtle changes in small patches of visual cortex in individuals (Glezer & Riesenhuber, 2013). Here, we looked at children ages 7-12, of various reading ability, and defined regions of interest in individuals’ native space, in order to examine the relationship between word selectivity and reading skill. Our findings suggest that over the process of learning to read, the VWFA becomes increasingly fine-tuned for words, and that word selectivity in VOTC is an essential component of skilled reading.

Declarations of conflicts of interest: none

## Acknowledgements

This work was funded by NSF/BSF BCS #1551330 to JDY, and Washington Research Foundation Funds for Innovation in Neuroengineering to ECK. We would like to thank the families who participated in the study, Patrick Donnelly for coordinating the study and help with data collection, and Deborah Burke for helpful comments on the manuscript.

---

**Supplementary Figure 1.**
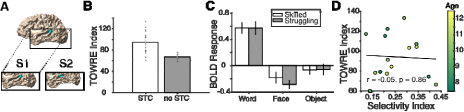
Selectivity in superior temporal cortex (STC) is not correlated with reading skill. (A) STC ROIs defined in individual subjects using a word > face contrast (B) Those with a STC ROI (n = 15) are significantly better readers than those without a STC ROI (n = 5) (t(18) = 2.66, p = 0.01). (C) Both skilled and struggling readers show high response to words in STC, and no response to other visual stimuli. (D) Word selectivity in STC is not correlated with reading skill.

